# Co-mutation based genetic networks to infer temporal mutation dynamics in ancient human mitochondrial genomes

**DOI:** 10.1101/2025.05.02.651746

**Authors:** Rahul K Verma, Jesualdo A Fuentes-González, Jason Pienaar, Janna Fierst

## Abstract

The evolution of modern humans is as fascinating as it is complicated. Various environmental and socio-political factors have played a significant role in shaping different migratory paths and lifestyles in ancient times and the establishment of current sub-populations around the globe. These factors have impacted the molecular evolution of mitochondrial DNA (mtDNA). To account for the role of genetic background and variable sites in shaping the migratory and evolutionary paths, we analyzed the ancient mtDNA as spatiotemporal co-mutation networks. Haplogroup-based analysis of the variable sites suggests a transition from hunter-gatherer to agrarian lifestyle during the Copper-Bronze transition period. Furthermore, the genetic interaction networks showed that COX and CYB genes behaved diversely with time and that there is an interplay of the genetic interactions with NADH Dehydrogenase genes specific to each age. We performed a phylogeny (tree) based gene network analysis and examined polymorphism to divergence ratios to complement the co-mutation networks. The tree-based gene networks exhibit similar topology with reduced co-mutation among the genes. The more traditional polymorphism/divergence analysis indicated that the CYB gene has been under long-term purifying selection. In contrast, other genes (ATP6, COX, NADH hydrogenases) have undergone episodes of purifying selection corresponding to different ages. We believe this network-based exploratory study provides insights into early lifestyle transitions and haplogroup establishment of the current human population.

## 1 Introduction

Biological systems exhibit complex functions from molecular to organ-system levels. These systems can be represented as interconnected entities in the frame of biological networks [1]. Traditionally, a biological network consists of a type of biomolecule as nodes (proteins, genes, or metabolites) depending on the system and the interactions between them be it physical, chemical or functional [2]. With the boom in the availability of more and more biological data, there are newer ways of considering the nodes and connections in biological networks.

One such way is to construct the networks based on the variable sites of DNA sequences as co-mutation/co-occurrence networks [3]. In most of the human cells, there is a nuclear genome and there is a cytoplasmic genome which is mitochondrial DNA. This mitochondrial genome is smaller in size and present in high copy number than the nuclear genome, which makes it a favorite tool of biologists to study human evolution along with paleontology [4]. The fossil record suggests that archaic humans (*Homo erectus*) first walked on Africa between 2 to 1.8 million years ago and likely migrated out of Africa for the first time probably between 300,000 and 280,000 years ago [5–7]. Genetic studies provide further evidence of biogeographical history of humans, especially when DNA collected from the archaeological sites is coupled radiometric dating of the samples it was extracted from [8]. However, it has traditionally not been trivial to extract and purify the DNA samples from fossils due to (a) degradation of chemical composition of the DNA sequence, (b) minute quantities, and (c) organic contamination [9]. Recent advances however, have facilitated retrieval of much higher quality DNA sequences from archaeological sites [10]. Nonetheless, the majority of human ancient DNA (aDNA) studies have focused on mitochondrial DNA (mtDNA) since it is present in multiple copies per cell and is often the only genetic marker that can be recovered from poorly preserved museum specimens and fossil remains. Ancient mtDNA therefore plays a critical role in the reconstruction of human history, including early human migration and admixture events, due to its high copy number and smaller size [11].

The mitochondrial genome, as is the case with the nuclear genome, has acquired genetic adaptations to different environments which can shed light on temporal and biogeographical evolutionary processes [12]. The central role that mitochondrial genes play in metabolism furthermore, allows us to use ancient mtDNA variation to infer how the availability of nutrients has influenced human migration and subsequent evolution [13, 14].

Mitochondria is the site of essential metabolic processes necessary to generate energy from proteins, carbohydrates, and fats [15, 16]. The availability of food and its nutritional content has been strongly linked with human population diversity [17–21] and the mitochondrial gene expression patterns [22]. The protein-carbohydrate makeup of a human’s diet influences the function of mitochondrial complex I both qualitatively and quantitatively [23]. Recent studies have even shown that mtDNA variation can also have diet-dependent effects on longevity [24, 25].

Current human subpopulations can be classified by multiple mitochondrial haplogroups that represent the major branching points in human mitochondrial phylogenetic tree. These haplogroups are defined as sets of mutations in mitochondrial DNA inherited from a common ancestor that typically vary with geographic region [26, 27]. Studies of ancient mtDNA have also revealed temporal variation in prevalence of these haplogroups [28]. A critical distinction, that will help us to understand the impact of the early lifestyle changes on modern human settlement, is if shifts in haplogroup frequencies between the different ages of human evolution reflect selective effects of diet, driven in part by the transition to an agrarian lifestyle, or if they simply reflect the genetic drift due to effective population size fluctuations. Currently, haplogroup H is the most frequent globally and is believed to have originated in Southwest Asia nearly 25,000 years ago, descending from the HV clade with two defining synonymous mutations, G2706A (16srRNA) and T7028C (CO1) [29, 30]. On the

European continent, haplogroup U is the oldest maternal haplogroup, dating to pre-agricultural populations [31] where its dominance at that time coincided with the rise of early farmers in the Neolithic Age [32]. Haplogroup U descended from haplogroup R through the defining mutations A11467G (ND4), A12308G (tRNA-Leu), and G12372A (ND5) nearly 43,000 to 50,000 years ago [29]. The shift from dominance of haplogroup U in Europe to the currently dominating haplogroup K coincides with transitions from hunter-gatherer societies to the early farmers of ancient humans [33–35] but it is not known if this shift was driven by agriculture-mediated selection on human mitochondria.

In the past two decades, a high number of complete mtDNA assemblies have been organized and made available through the ancient human mitochondrial genome database (AmtDB) which contains complete information regarding the source, age and haplogroup of the samples [36]. Here, we use a combination of unconventional co-mutation network analysis, along with conventional phylogenetic reconstruction and within human polymorphism to human-chimp divergence ratio analysis to investigate patterns of mtDNA variation over the last ~10,000 years of human history. We constructed and analyzed co-variation-based genetic networks for ancient mtDNA for ~10,000 years, covering six ages of human evolution. Based on this analysis. We identified a set of prevalent genes for each age and found that NADH dehydrogenase complex genes interact with cytochrome B and cytochrome oxidase complex genes throughout each age, suggesting a functional role for these genes in defining the nutritional habits throughout the human evolution. The co-mutating mitochondrial genes in each age also clustered as per the temporal sequence of the ages suggesting a role of demographic factors. We furthermore constructed gene trees using a maximum likelihood (ML) algorithm for each age and upon further analysis, found that the phylogenetic concordance and discordance in these populations correlate with reduced/enhanced genetic interaction networks, respectively. Lastly, we determined the Dn to Ds ratios within humans per protein coding mitochondrial gene for each age (*ω*_*polymorphism*_) and the divergence Dn to Ds ratios through alignment to the Chimpanzee mitochondria (*ω*_*divergence*_) and used these to perform McDonald-Kreitman tests for selection that we interpreted with the neutrality index (NI = *ω*_*polymorphism*_ / *ω*_*divergence*_).

The number of samples considered in this study is shown. Abbreviation of the ages are shown in parenthesis.

## 2 Results and Discussion

To study the temporal dynamics of mtDNA haplogroups we downloaded all complete mtDNA FASTA sequences for which C14 dating was available from the Ancient mitochondrial DNA Database (AmtDB) [36]. We obtained 812 ancient mtDNA sequences sampled from six different ages, ranging from the Mesolithic Age to the Middle Age (Table 1) that we were able to align to rCRS along their full lengths.

**Table 1.** Number of samples and the period of each age.

| Age | Period range | Sample size |
| --- | --- | --- |
| *Mesolithic(Meso)* | 9755 BCE to 1850 BCE | 82 |
| *Neolithic(Neo)* | 8250 BCE to 1519 BCE | 206 |
| *Copper(Cop)* | 4725 BCE to 1690 BCE | 108 |
| *Bronze(Bro)* | 4452 BCE to 230 BCE | 268 |
| *Iron(Iro)* | 1390 BCE to 615 CE | 76 |
| *Middle(Mid)* | 10 CE to 1264 CE | 72 |

### 2.1 Quantifying Variable Sites

We built the co-mutation networks after extracting the variable sites for each age as discussed in the Methods. We found that these networks were disconnected which means that there were some pairs of variable sites that were not co-mutating. Moving forward, we considered only that component of the network which consisted of maximum pairs of variable sites henceforth called the largest connected component for further analysis of all networks. The variable positions for each age were mapped with the ancestral SNPs from Mitomap [37]. We found that the Mesolithic Age had the highest percentage (6.8% of 219 variable sites) and the Middle Age had the lowest percentage (1.0% of 560 variable sites) of ancestral SNPs. The high proportion of ancestral variants suggests that older populations have undergone fewer bottlenecks and admixture events with more basal haplotypes. In contrast, the lower percentage of ancestral SNPs in recent ages suggests genetic drift and population expansions. This could be attributed to the transition from foraging to farming lifestyle, since agriculture provided basis for bigger population sizes and admixture [38].

The global structural properties [39] of all the networks are shown in Table 2. We can see from the table that not all of the significantly co-mutating variable sites were participating in the network. To quantify the contribution of variable sites of individual genes in each network, we plotted the ratio of the number of total variable sites in each gene to the variable sites forming that network (Fig 3). We observed that the tendency of variable sites to co-mutate increased (fewer genes with non-co-mutating sites) from the Mesolithic Age to the Middle Age. This suggests that the sub-populations have become more homogenized probably with the advent of an agrarian lifestyle [40, 41]. Such a lifestyle supports more extensive gene flow, reduced heterogeneity, and promotes greater coherence in mutation patterns across the populations [42]. Interestingly, the smallest gene ATP8 is present in three different ages and the largest gene ND5 is present in four ages along with other genes. Along with this, the ND4L gene, another small gene with only 98aa long protein, had almost 50% of its variable sites not participating in the network in the Mesolithic Age and then only observed during the Middle Age where less than 10% of variable sites did not participate in the network. This suggests that the participation of variable sites in the network’s formation is independent of the gene size. Furthermore, the networks are sparse which reflects the specific nature of interactions between the variants. There was a negative correlation between node degree and the clustering coefficient in all the ages except for Copper and Middle Ages (Fig 4). In the Bronze Age, this correlation is highly prominent and suggests extensive hierarchy in the network. This could imply that the advancement of technological development during the Bronze Age with the introduction of global trade, dietary alterations because of agriculture, and subpopulation interactions led to higher population densities [43]. In addition, the presence of high modularity suggests that the networks are organized in tightly linked clusters or locally dense regions where nodes within the cluster are more likely connected to each other than with nodes in other clusters. This grouping of the variable sites as clusters points to the functional or evolutionary constraints in the mitochondrial genome irrespective of the time periods.

**Table 2.** Structural properties of co-mutation networks.

| Age | Variable Sites | $N_{LCC}$ | $E_{LCC}$ | $\langle k \rangle$ | $\langle cc \rangle$ | Modularity |
| --- | --- | --- | --- | --- | --- | --- |
| *Mesolithic* | 219 | 185 | 1087 | 11 | 0.74 | 0.67 |
| *Neolithic* | 530 | 511 | 3400 | 13 | 0.67 | 0.71 |
| *Copper* | 491 | 461 | 3005 | 13 | 0.76 | 0.82 |
| *Bronze* | 722 | 712 | 4003 | 11 | 0.26 | 0.59 |
| *Iron* | 375 | 335 | 1723 | 10 | 0.80 | 0.83 |
| *Middle* | 580 | 560 | 6338 | 22 | 0.84 | 0.79 |

Variable sites are number of nodes participating in network construction at $P_{ij} < 0.05$ , $N_{LCC}$ is the number of nodes in largest connected component, $E_{LCC}$ is the number of connections, $\langle k \rangle$ is the average degree and $\langle cc \rangle$ depicts the average clustering coefficient of the largest connected component.

**Fig 1.**
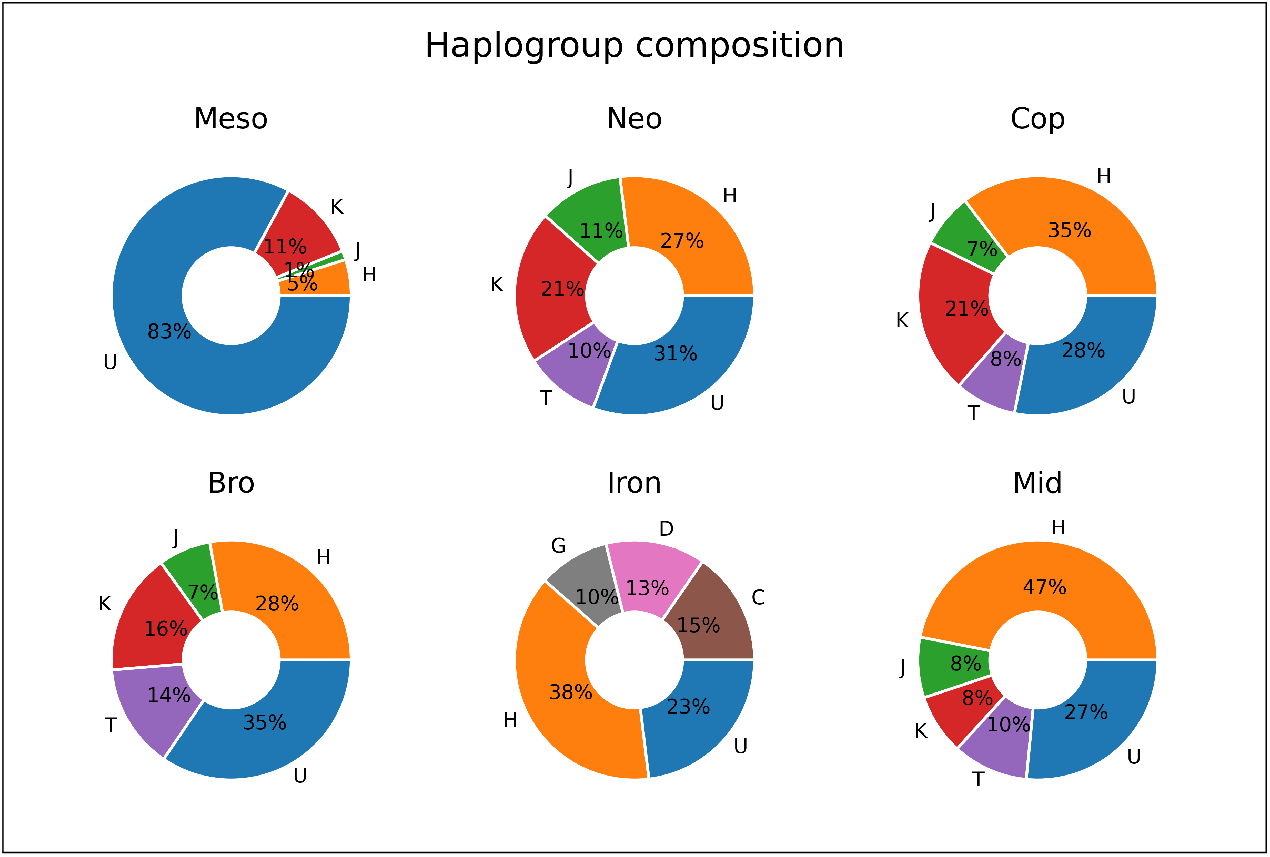
The haplogroup composition of mitochondrial samples in each age. Only major haplogroups are considered here.

**Fig 2.**
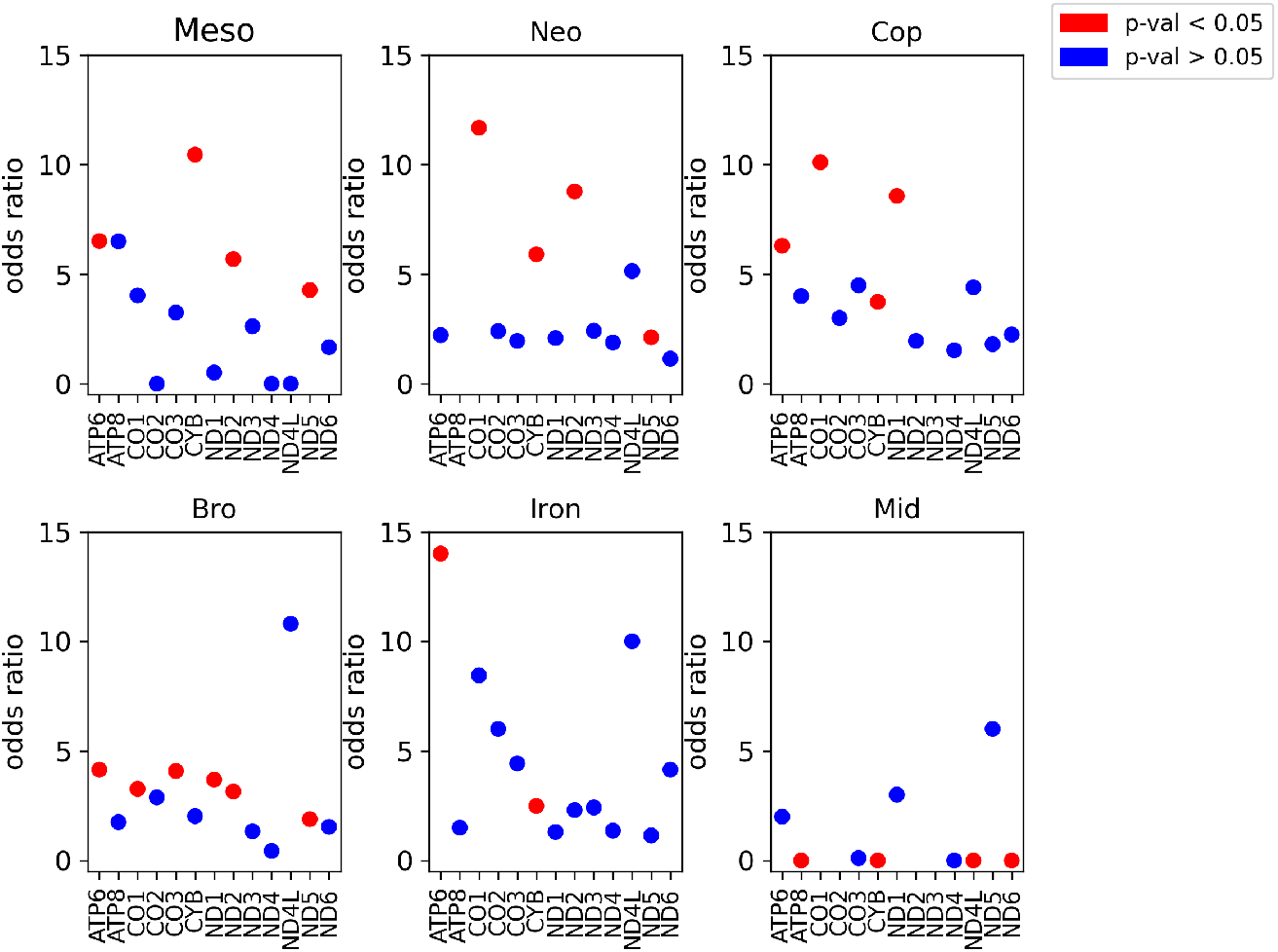
MK test is performed to calculate the odds ratio between archaic human population and chimpanzee to get the neutrality index.

**Fig 3.**
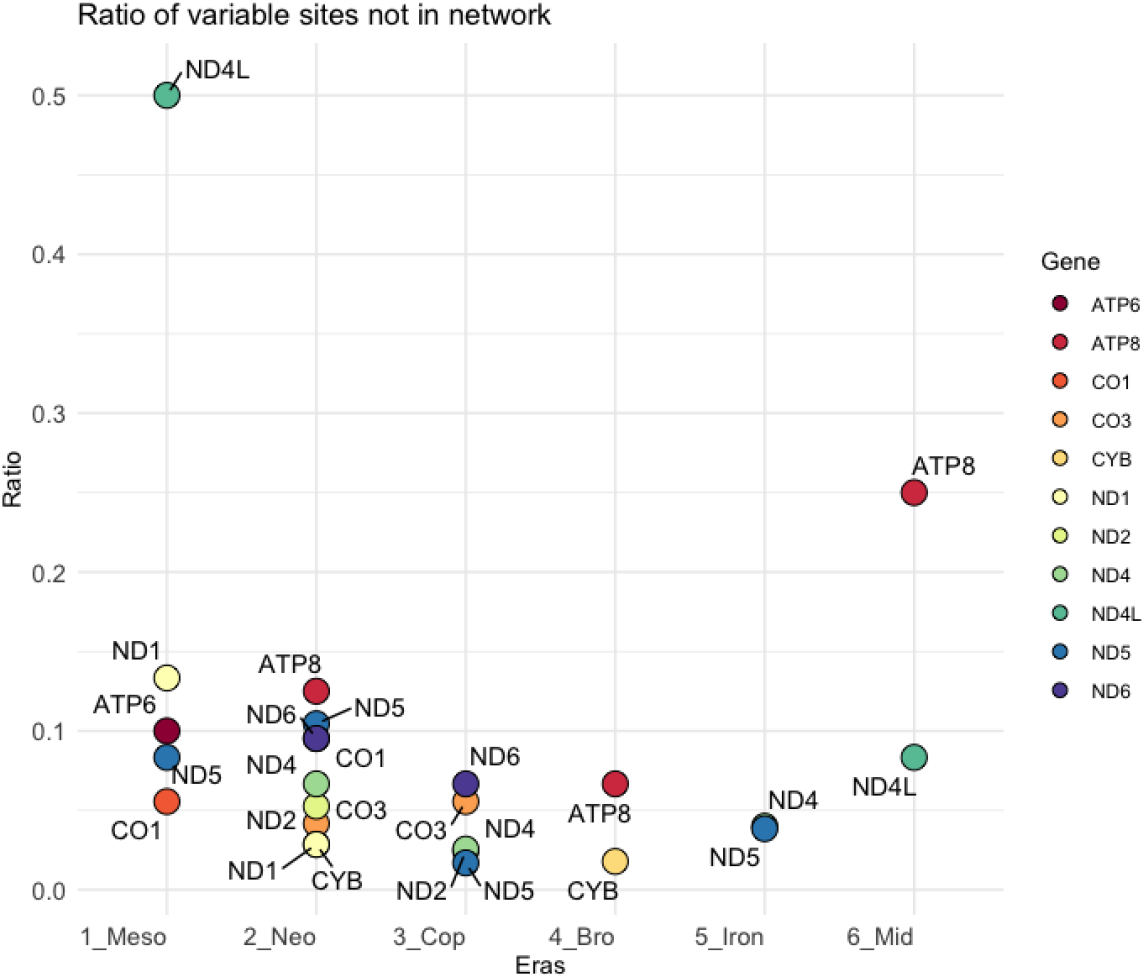
The ratio of total variable sites to the variable sites taking part in the network construction.

**Fig 4.**
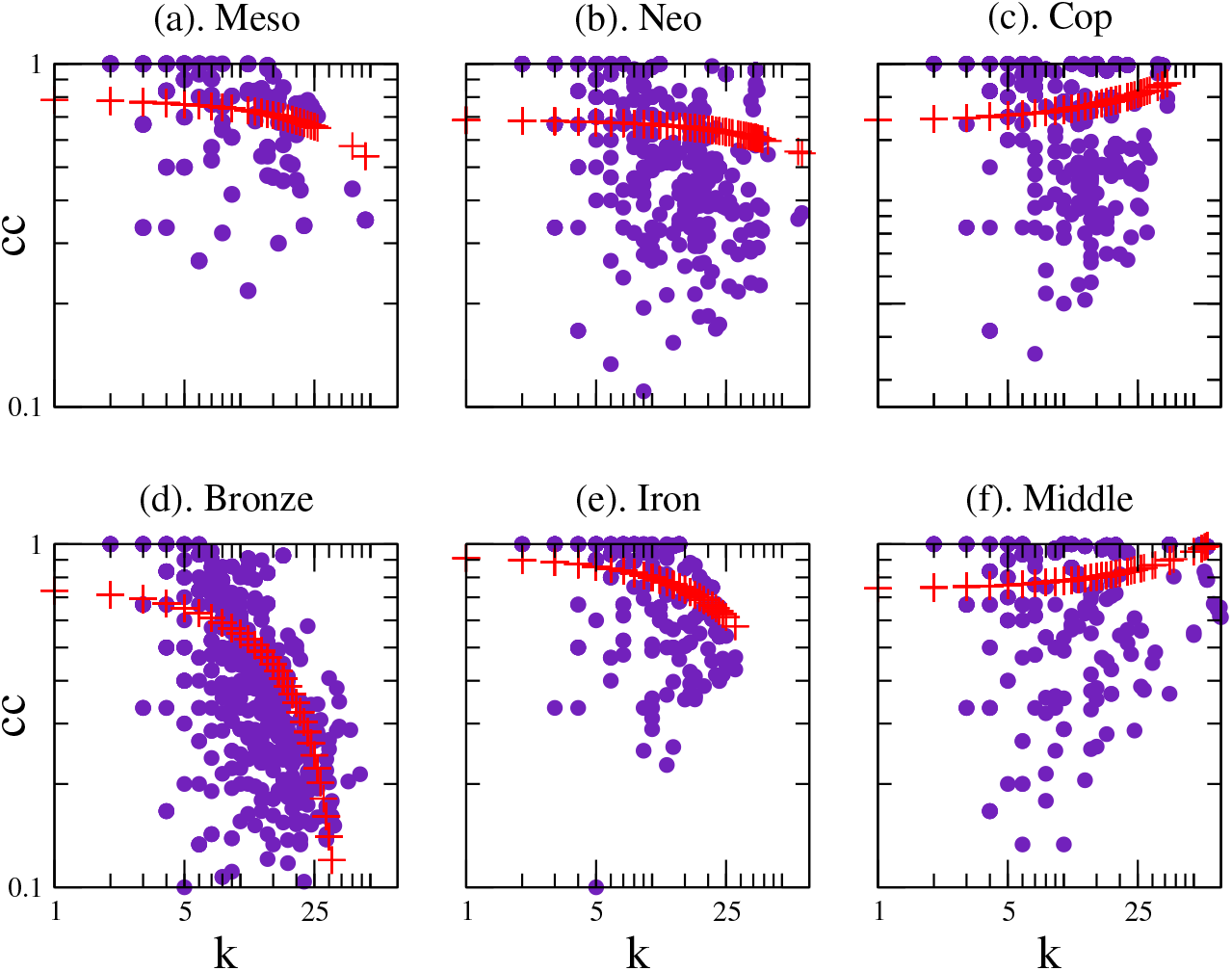
The relation between the node (variable sites) degree (k) and clustering coefficient (cc) is plotted here for each era. The plots show an inverse relationship except for Copper and Middle Age. In the Bronze Age, the scale-free nature is quite visible.

#### Evolutionary dynamics of hub variants across the eras

High-degree nodes or ‘hub odes’ are of particular importance as they are the nodes with extensive connections that makes the network stable and robust. These nodes are likely to represent critical variable sites, possibly indicating evolutionary hot spots that drive the interaction patterns within the network. Next, we discuss such variable sites in each age, along with the additional information that the MK (McDonald–Kreitman) test and neutrality index (NI = *ω*_*polymorphism*_ / *ω*_*divergence*_) tell us about the genes these nodes are found in (Fig 2)

*Mesolithic Age:* Position 9477 of the CO3 gene is hub node which is a defining variant for the U5 haplogroup and is a non-synonymous variant causing a change from valine to isoleucine at position 91 in CO3 protein subunit. This variant is responsible for high sperm mobility and quality in human males [44]. Despite the NI for the CO3 gene of 3 (Fig 2) suggesting functional constraints on this gene in the Mesolithic Age, the MK test did not detect a significant departure form neutrality, which is consistent with valine and isoleucine being interchangeable as non-polar amino acids in the transmembrane portion of the protein.

##### Neolithic Age

Position 9428 of the CO3 gene is a hub node which is a synonymous variant implying its influence may lie in mRNA stability, codon bias, or nascent protein folding [45]. The NI for CO3 of 2, again implies possible purifying selection on CO3 in the Neolithic, but again, it was not a significant departure from neutrality.

*Copper Age:* This period is marked by the diversification of hub nodes with variants 6116 and 6047 in the CO1 gene and 15511 in CYB gene as critical hubs. Among the three hub nodes, the 6047 in the CO1 gene is a marker of the U4 haplogroup. The presence of multiple hub nodes may indicate complex selective pressures during this period, possibly reflecting a period of migration and genetic admixture [46]. The association of haplogroup markers with hub nodes underscores the dual role of these sites in both defining genetic lineages and contributing to mitochondrial function. Interestingly, both the CO1 and CYB genes are significant for the MK test and the respective NI’s of 10 and 9, indicate strong purifying selection on both these genes in this period. Looking back in time, the CO1 gene was also under strong purifying selection in the Neolithic Age, whereas the CYB gene was constantly under strong purifying selection. Looking forward, the CO1 gene is under constant purifying selection in the Bronze and Iron Ages whereas the CYB gene appears to be neutral in the Bronze Age (but with a NI ~ 2), again under purifying selection in the Copper Age. Both genes for the most part appear to have been under purifying selection across the periods examined here.

#### Bronze and Iron Ages

Position 930 in the 12s rRNA gene is a hub node. 12s-rRNA is part of the ribosomal machinery suggesting the influence on the translational regulation of mitochondrial genes. In the Iron Age, variant 12705 of the ND5 gene is a hub node which is again a synonymous variant and the MK test suggests it is evolving neutrally in the Iron age but under purifying selection in the Bronze (as well as in the Mesolithic and Neolithic Ages).

#### Middle Age

Position 15454 of CYB gene is hub node which is a synonymous variant and a marker of the U3 haplogroup.

Mostly, the hub nodes are synonymous variants which suggests consistency in metabolic homeostasis. CYB, as mentioned above, consistently turns out significant for the MK test and the NI*>*1 suggests long-term, strong purifying selection throughout the ages.

Overall, the genes containing Hub nodes were enriched for significant and strong purifying selection where the non-Hub containing genes are enriched for neutrality. The ATP6 gene is an exception in that it does not contain any hub node variants, but the MK tests for this gene where significant in the Mesolithic, Copper, Bronze and Iron Ages and the minimum NI was 4 indicating consistent, strong purifying selection (Fig 2). The ATP6 gene is involved in the ATP production and is the last complex of the electron transport chain. Studies showed that ATP6 has undergone purifying selection and constrained the amino acid changes to maintian the fuctional integrity of this gene [13, 26].

Next, we checked for the haplogroup composition of each period which was available as metadata at the AmtDB (Fig. 1). We analyzed haplogroup composition across the periods to get a comprehensive view on the genetic and cultural dynamics of contemporary populations. This analysis showed that the Mesolithic age was completely dominated by haplogroup U, while in the Neolithic and Copper Ages haplogroups U, H, and K were dominant.

Haplogroup U found to be associated with the hunter-gatherer group and haplogroup H was found to be associated with the farmer group [35, 47]. The dominance of the haplogroup U in the Mesolithic Age reflects its strong association with the hunter-gatherer lifestyle. This observation aligns with the broader understanding that Mesolithic people were heavily dependent on foraging and hunting. Following the Mesolithic Age, there was a shift in the haplogroup composition with the emergence of haplogroups H and K alongside haplogroup U. This pattern is consistent with the introduction of farming and development of the agricultural practices in Europe. The presence of haplogroup K further supports the sedentary farming lifestyles [48, 49]. In the Copper, Iron, and the Middle Ages, the haplogroup H was found to be dominant, which supports the idea of agricultural developments during these periods. In the Bronze Age, haplogroup U underwent extensive range extension and substitutions due to genetic drift [50]. This suggests that the presence of haplogroup U was majorly shaped by population bottlenecks or migration events.

### 2.2 Temporal dynamics of mtDNA haplogroups

Over 80% of the samples from the Mesolithic age belong to haplogroup U along with haplogroups K, H and J (Fig. 1). Haplogroup H was present in the Mesolithic Age at a low frequency of 4% of the samples. During the Neolithic Age haplogroup U remained the most frequent. Haplogroup U comprised 27% of the samples with an additional 10 haplogroups identified at different frequencies. mtDNA haplogroup diversification continued with 14 major haplogroups identified in the samples from the Middle Age. Haplogroup U was found to be the dominant mtDNA haplogroup during the Mesolithic, Neolithic, and Bronze Ages while Haplogroup H was dominant during the Copper, Iron and the Middle ages. The pattern of haplogroups observed is in line with the lifestyle of the populations.

We discussed the pattern of the H and U haplogroups in our networks from the Mesolithic age to the Middle age in previous section. Now, we were interested in the dynamical pattern of co-mutation partners of the variable sites of these haplogroups. We observed that the co-mutations of haplogroup H were dominantly belonging to L, M and H haplogroups, and that of U haplogroups were dominantly belonging to L, M, H and U haplogroups. L being the root mitochondrial haplogroup [6], it is expected to observe such interaction patterns. M haplogroup evolved parallel to the N haplogroup, parent clade of H and U haplogroups. The co-mutations of H and U haplogroups variable sites belonging to M haplogroups suggest possible interactions at the population level due to geographical vicinity. On further observing the first level clade of these haplogroups, it was noticed that the H haplogroup interacts prominently with the variable sites of the H1 clade up to Iron age and the L3 clade in the Middle Age. In case of the U haplogroup, the most prominent clade was U5 for the Mesolithic, Neolithic and Iron Age, and H1 clade for the Copper, Bronze and Middle Ages. This suggests that the H haplogroup has evolved closely within the population and being restricted to interaction with people of similar lifestyles. We can say that the group of people who transitioned to the agrarian lifestyle contained themselves to a specific region and interacted more often with people having similar lifestyles.

#### 2.2.1 Global nodes

Since we were interested in the overall dynamics of variable sites across ages, we found 25 global (common) nodes that were present in all the networks. The global nodes provided us with a common reference to derive the overall changes in the gene interactions across the ages. We mapped the first neighbors of these global nodes to their corresponding genes to look into the dynamics of the genes across all the ages (Fig. 5). We considered 5 protein-coding genes CO1, CO3, CYB, ND2, and ND4 due to the variability in their degrees across the eras. The degrees in the Neolithic Age were observed to be highly varied and spread across a long range. There was a drop in the degree of the genes in the Copper Age which was maintained as is for following periods except for the CYB gene. The Copper Age seems to be a point of transition from hunter-gatherers to farmer lifestyle as seen from the haplogroup analysis (Fig. 1. Similar behavior was observed here in terms of the degree of the genes co-mutating with common variable sites. This suggests that the dynamics of these genes synchronized as the populations transition to a unified lifestyle. One peculiar behavior of the CYB gene was that its degree drastically increased after the Copper age.

**Fig 5.**
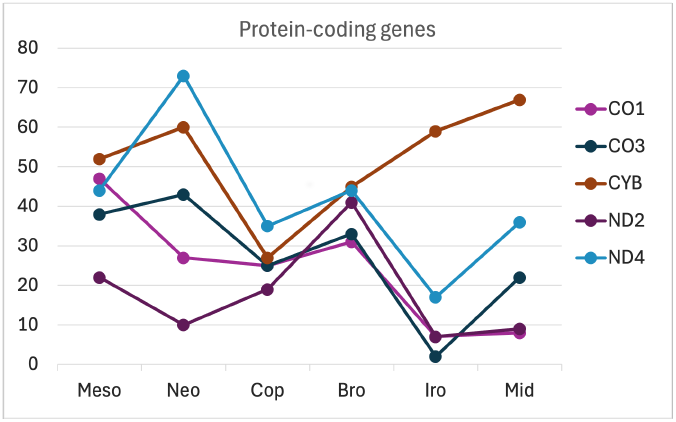
The degree of the genes of first neighbors of common nodes.

## 3 Genetic interaction networks

The gene-pair weights were calculated as the sum of the number of times each pair was present in the network. Then, we normalized the weights by the sum of the length of genes for any given pair of genes and scaled by the maximum weight. We compared the protein-coding genes with the corresponding random networks for each age and presented only the genetic interactions (Genetic-enhanced) which were either one standard deviation higher than the random one or one standard deviation lower than the random ones (Genetic-reduced). We found that the different genes showed significant interaction weights in the different periods (Fig 6). It was observed that, in all the periods, more interactions were showing lesser weight as compared to interactions showing higher weight than the random ones. However, it is difficult to interpret these reduced and enhanced interactions in terms of mitochondrial biology, we try to focus on dominant interactions in general for all the periods. We will use the enhanced and reduced interactions together to discuss the results. As we know that the co-mutation interactions between genes can provide the functional and evolutionary aspects of the genome. These interactions also highlight epistatic relationships, where the combined effect of mutations differs from the sum of their individual effects, often shaped by purifying or compensatory selection. The gene-gene interactions can often modulate evolution at the molecular level by inhibiting or promoting specific mutation combinations [51]. In terms of co-mutation, the edge weight represents the tendency of co-occurrence of variations between genes. Now, we look at the more prominent interactions in each age, in Mesolithic Age, ND5 with ND4 and CYB with ND4 genes; in Neolithic Age, ND1 with CO3 and CYB genes, and ND5 with CO1, CYB, ND1 and ND6 genes; in the Copper Age, CO1 gene with ND6, and ND2 with ND5 and CO1 genes; in

**Fig 6.**
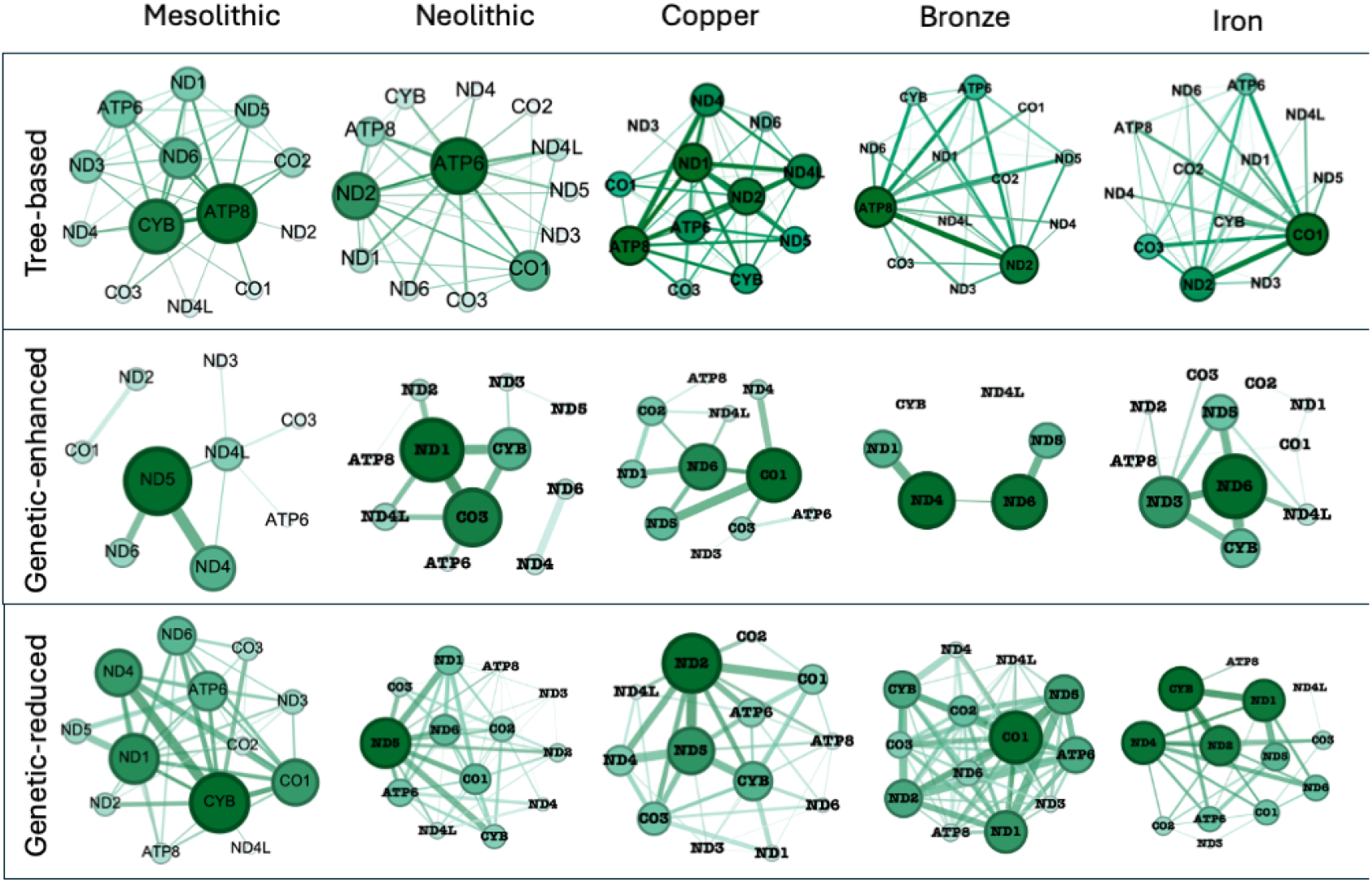
The tree-based gene networks and the genetic interaction networks are compared here. The Genentic-reduced are the genetic interaction networks that show co-mutation lesser than the corresponding random networks while the Genetic-enhanced show higher co-mutation than the corresponding random ones.

Bronze Age, ND4 with ND1, ND6 with ND5, and CO1 with multiple genes; in Iron Age, ND6 gene with ND3, CYB with ND5, and CYB, ND4, ND1, ND2 genes forming prominently high weight interactions. As we can see, in the enhanced genetic interactions, ND genes are more prominent along with CO genes in the Neolithic and Copper Ages. Cytochrome c oxidase (CO complex) is the last complex in the electron transport chain which regulates mitochondrial respiration. This is the point where oxygen is consumed and the protons are pumped into inter-membrane space, so, this is a crucial point where ATP and ROS productions are regulated [52].

### 3.1 Tree-based gene networks

To complement the co-mutation networks we used a conventional method to implement the phylogeny. We constructed thirteen phylogeny trees of protein-coding genes for each age. We started by contrasting several nucleotide substitution models to identify which ones better described the protein-coding gene alignments for each age. The networks with reduced co-mutation followed relatively similar topology as the tree-based gene networks, as discussed further. We compared the degrees of the genes of tree-based networks and that of the genetic networks. The tree-based networks showed a similar trend as that of the genetic networks with reduced co-mutation. This suggests that the genetic networks with reduced co-mutation are informative of the shared ancestry while the genetic networks with a high co-mutation might be capturing functional aspects of the mitochondrial genome in terms of temporal and lifestyle-based evolution [53, 54]. We also calculated the correlation distance between the periods based on the genes from the genetic interaction networks and plotted them as a dendrogram (Fig. 7). The dendrogram of genes clearly makes the separate clusters for periods with a branching pattern that reflects temporal associations. The first branching clusters the periods into two groups one with the Mesolithic and Neolithic Ages and the other with the remaining four periods. Iron and Middle Ages along with the Copper and Bronze Ages formed two separate clusters. The Middle and Iron Ages were farthest from Mesolithic Age which is in line with their temporal state. In the dendrogram of genes the Copper and Bronze Ages were closest to each other. The tree of genes-based similarity captures the temporal dynamics.

**Fig 7.**
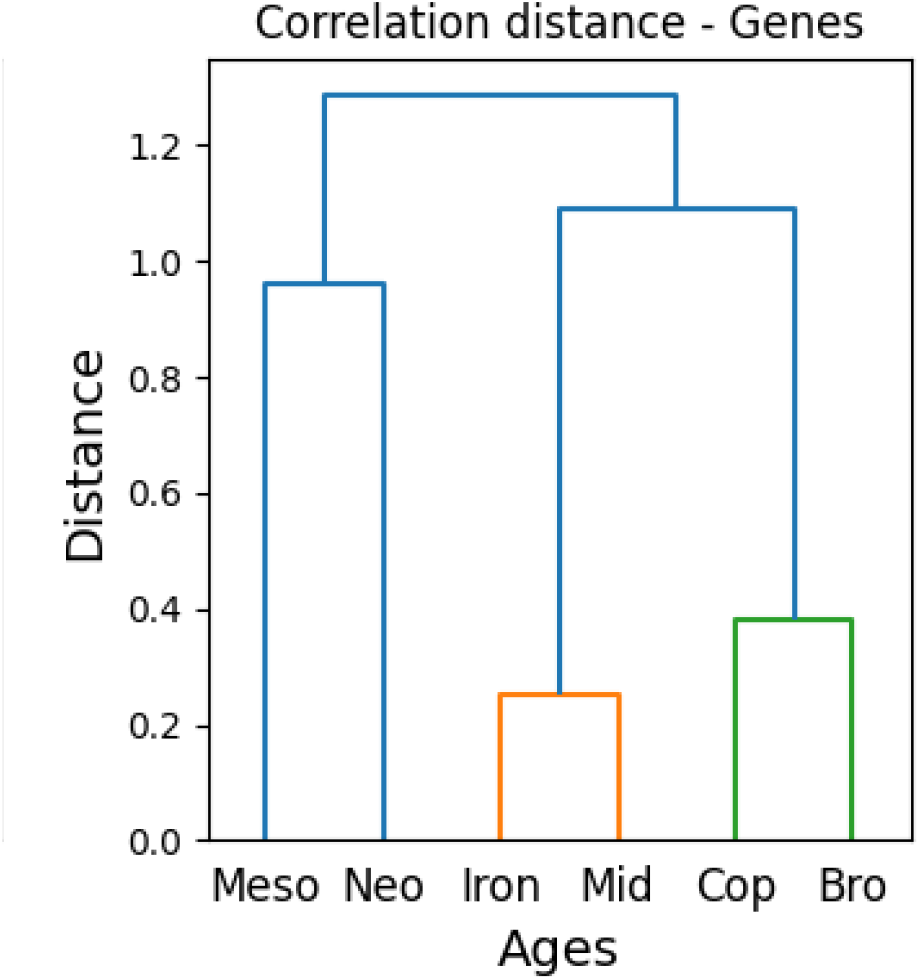
Jaccard’s distances between the eras were calculated based on presence of genes and plotted as clusters using the correlation metrics.

**Fig 8.**
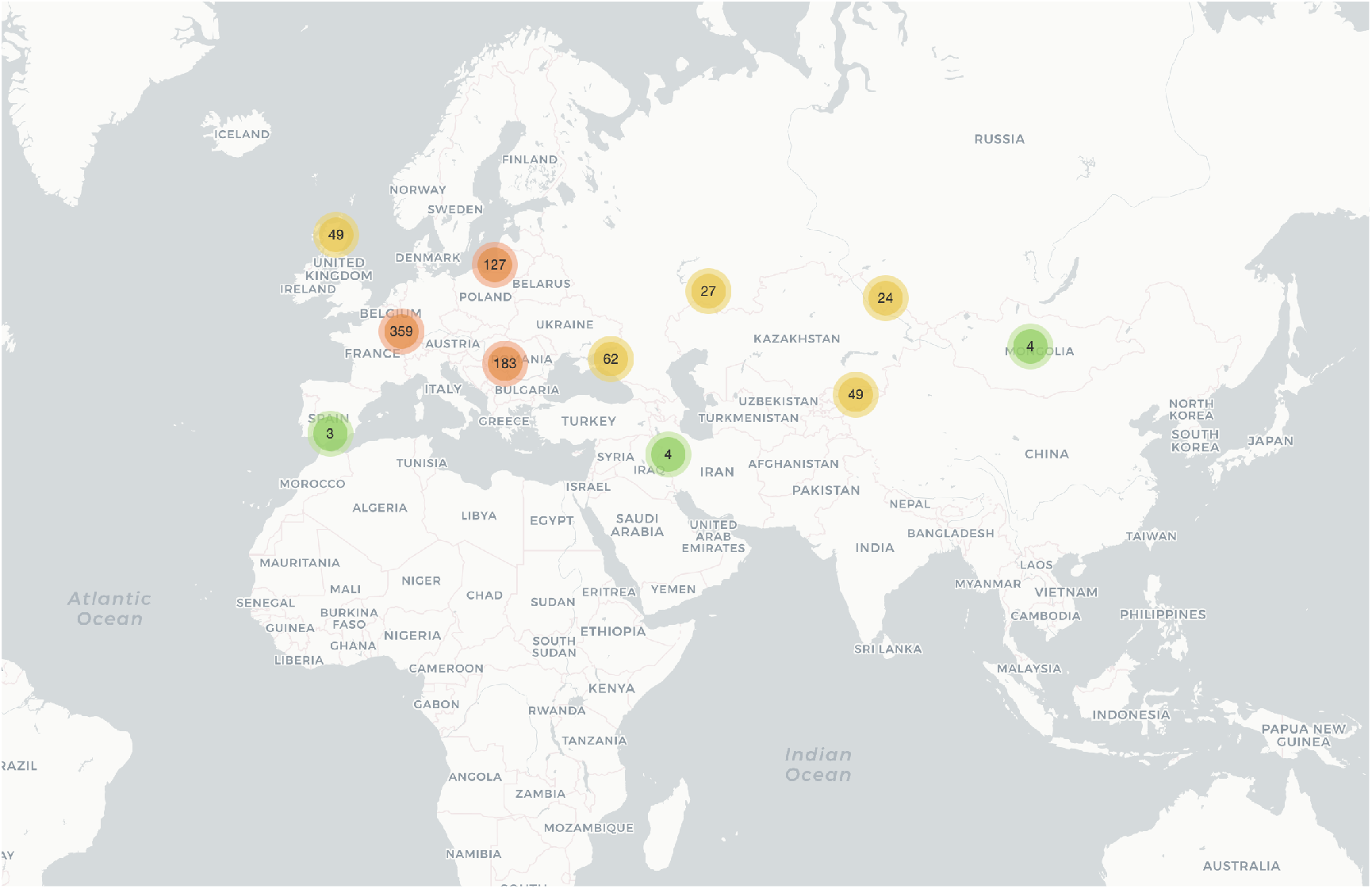
Geographical locations of the samples used in this study downloaded from AmtDB.

In summary, the complex temporal dynamics of variations in the mitochondrial genome is readily captured by the co-mutation networks. The statistical analysis of these networks provides a better understanding of the genetic interactions for each age. The presence of CO1 and CO3 genes as the high degree nodes in all ages suggests for the potential role of cytochrome oxidase complex in diet (lifestyle) based evolution. Additionally, as we go from the Mesolithic Age to the Middle Age, it is observed that the tendency of the variable sites to pair with other variable sites increases, which further strengthens the idea of a homogeneous population as a societal structure that is most likely based on the agrarian lifestyle. As it is known the network analysis is an established approach to analyze the complex systems, here, we complemented it through another established approach of phylogenetic (tree-based) analysis. We found that these tree-based genetic networks identify the similar set of genes as found in the reduced co-mutation networks which suggests the presence of both concordance and discordance in the evolution of the lifestyle. In other words, the the reduced co-mutation networks are representing the phylogeny based evolution while the increased co-mutation networks captures the functional aspect of the genome in term of co-mutation. Overall, the networks analysis being a powerful tool if, combined with modified traditional approaches can provide a better insights into complex dynamics of genome evolution.

## 4 Materials and methods

### 4.1 Data acquisition and pre-processing

Complete mtDNA FASTA sequences were downloaded from the Ancient Mitochondrial Database (AmtDB) [36] for which C14 dating was available. We grouped these sequences according to the available metadata for each Age (Table 1). We aligned all the sequences using Clustal Omega with default parameters. Any sequences with insertions/deletions were removed so as to keep the length of aligned sequences the same as the human mitochondrial reference sequence (rCRS).

### 4.2 Co-mutation Network Construction

**Step 1:** Any position having more than one variant present at least twice within the samples was considered a variable site. These variable sites were extracted from the aligned sequences for each Age separately for constructing co-mutation networks. For genomic equality, ambiguous nucleotides represented as ‘X’, ‘M’, ‘Y’, etc. were replaced with ‘N’ for all the sequences. Since these sequences were extracted from fossil remains, there were various positions where nucleotide is unknown and denoted as ‘N’. Therefore, we incorporated the presence of an unknown nucleotide for considering a position as a variable site with the threshold of 5%, which was set empirically. For example, for variable site positions where ‘N’ was present *<* 5%, we still considered it as a variable site but did not include the sequences carrying ‘N’ for further calculations.

**Step 2:** For the construction of a network, a node was represented by the position of a variable site, and the edge was represented by co-mutation frequency between any two nodes. We defined co-mutation frequency for a pair of variable sites, *C*_(*ij*)_ as,

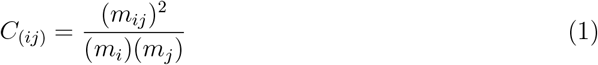

where *m*_*ij*_ is number of times minor alleles occur together (defined as co-mutation pair) at *i*^*th*^ and *j*^*th*^ positions, respectively, *m*_*i*_ is the total number of times minor allele occurs at *i*^*th*^ position and *m*_*j*_ is total number of times minor allele occurs at *j*^*th*^ positions individually.

**Step 3:** For checking the significance of any co-mutation pair, we defined and calculated a *P*_*ij*_ such as,

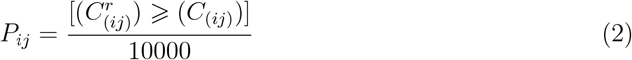

where 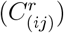 is the number of times two nodes co-mutate randomly, the total number of random realizations was set to 10,000. To generate random simulations, 10,000 permutations were performed for nucleotides at each variable position and we considered only those realizations where co-mutation occurred more often than the real one. Further, we took only those co-mutation pairs where the *P*_*ij*_ is *<* 0.05.

**Step 4:** Nodes were mapped to their corresponding genes for each co-mutation network to achieve one weighted gene-gene interaction network for each Age. Since two or more co-mutation pairs may belong to the same gene or a pair of genes, each link was counted as many times as it was found, and this number was considered as weight for that pair of genes. For example, the co-mutation pairs (3352-7623) and (4125-8054) would map to *ND1-CO2* gene-pair, so this edge was counted twice, and so on, similarly, the co-mutation pairs (3352-3489) would map to a self-loop for *ND1* gene. Since the variable sites contributed by each gene were in proportion to their lengths (except *Control region*), the weight of each gene pair was normalized by the sum of the total length of both the genes for that gene pair. Additionally, we constructed gene-gene interaction (GGI) networks using the co-mutations of the (i) common nodes and (ii) exclusive nodes for all three regions. For (i), we took the nodes commonly present in all the networks and scanned for their co-mutating partners, and mapped these co-mutating nodes to respective genes. Similarly, exclusive nodes for each region were used to construct the corresponding GGI network.

### 4.3 Phylogeny-based gene networks

In order to identify whether co-mutation interactions (with either high or low weights) were reflecting patterns of common ancestry, we contrasted them with analogous networks but informative of phylogenetic congruence by constructing thirteen gene trees of protein-coding genes for each age. The models incorporated various combinations of nucleotide substitution frequencies and rates, fraction of invariant sites, and rate heterogeneity under a gamma distribution with five categories [55]. Model fit was assessed under Maximum Likelihood (ML) and the candidates were compared under the Bayesian Information Criterion [56]. The preferred candidates from this comparison (i.e. those reporting the lowest scores) were used as substitution models to infer gene trees for each Age under ML [57]. We conducted a heuristic search using Neighbor Joining as the initial tree which was subsequently rearranged under Subtree-Pruning-Regrafting. All these evolutionary analyses (i.e. model comparison and phylogenetic inference) were conducted in MEGA11 [58, 59]. The resulting gene trees were used to construct networks of protein-coding genes for each period. To do this, we computed weighted Robinson-Foulds distances [60] among all gene trees for each period using the RStudio (4.3.1) package phangorn v2.11.1 [61]. The weighted Robinson-Foulds distance accounts for both tree topology and branch lengths, which allowed us to contrast the co-mutation networks not only in terms of tree shape but also in terms of accumulated molecular diversification. It is a commonly used distance metric [62] that has shown good performance under similar phylogenies [63], which is expected for inferred gene trees that are based on the same individuals and time bins.

### 4.4 Jaccard distances

We calculated the Jaccard distances between the ages based on the variable sites and corresponding genes from the co-mutation networks.

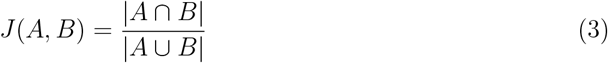

where, *J* (*A, B*) is Jaccard similarity coefficient, and A and B are sets of genes for a possible pair of genes.

